# Organelle scaling over a 100-fold cell size range

**DOI:** 10.64898/2026.05.13.724986

**Authors:** Alison C.E. Wirshing, Daniel J. Lew

**Affiliations:** Department of Biology, Massachusetts Institute of Technology, Cambridge, MA 02139

## Abstract

Cell size in a proliferating cell population generally varies over a limited range (∼2-4-fold). Within such populations, organelle content increases with cell size maintaining a relatively constant organelle density (amount per cell volume). However, cells of different types can differ greatly in cell size as well as in organelle composition. In such cases, it is often unclear to what degree, if any, the differences in organelle composition are due to the difference in cell size. In principle, this issue could be resolved by examining situations where a proliferating population of cells of the same cell type exhibit much greater size variation. Here we characterize how organelle content scales with cell volume in the polymorphic fungus, *A. pullulans*, whose proliferating cells span a ∼100-fold size range. We find that mitochondria and ER content increases in proportion to cell volume, while this is not the case for vacuoles and peroxisomes. Thus, organelle composition is affected by cell size in this system.

## INTRODUCTION

Cell size is a fundamental property that affects many aspects of cell function. For most proliferating cells, cell size is maintained within narrow limits by mechanisms that coordinate cell cycle progression and growth as a function of size (Campos et al., 2014; Di Talia et al., 2007; Sveiczer et al., 1996; Wang et al., 2000). Such cells generally double in size and then divide in half. However, other cells adopt different patterns of growth and division. Some hyphal fungi can grow dramatically from spores without dividing, and then generate hundreds of new spores (Heitman et al., 2007). Similarly, there are bacteria (Chimileski et al., 2024; Pląskowska et al., 2023), algae (Ivanov et al., 2019), chytrids (Medina et al., 2025), protists (Francia and Striepen, 2014), and other cells that alternate between periods of uninterrupted growth and periods of multiple division. These proliferative strategies can lead to much larger size differences among cells of a similar type. For cells of very different size, it is unclear how the demands for different organellar functions might scale with cell size.

Cell volume and surface area both change as cells get bigger, but the relation between them depends on the cell’s geometry. A cylindrical cell like a hypha growing at one end would increase both its volume and its surface area in parallel, but a spherical or ovoid cell would increase its volume faster than its area. For such cells, the changing surface-to-volume ratio may necessitate adjustments in the relative amounts of internal organelles. Here, we document the scaling behavior of various organelles in the black yeast *Aureobasidium pullulans*, whose approximately ovoid cells span an unusually large ∼100-fold size range.

*A. pullulans* is a poly-extremotolerant fungus that thrives in an extraordinarily diverse range of habitats, with varying salinity, pH, temperature, and nutrient availability (Adams et al., 2013; Ademakinwa and Agboola, 2016; Andrews et al., 2002; Bennamoun et al., 2016; Branda et al., 2010; Coleine et al., 2021; Grube et al., 2011; Gunde-Cimerman et al., 2000; Humphries et al., 2017; Nonnenmann et al., 2012; Olstorpe et al., 2010; Wang et al., 2009). *A. pullulans* has both hyphal and budding yeast forms (Goshima, 2022; Ramos and García Acha, 1975). In its yeast form, mother cells can contain variable numbers of nuclei, and they produce different numbers of buds in a single cell cycle by multi-budding (Mitchison-Field et al., 2019; Wirshing et al., 2025, 2024). Buds generally inherit only one or two nuclei, and then produce only one or two buds (Petrucco et al., 2025). However, they can also forgo budding and engage in “nuclear cycles” that increase the cell’s volume and double the number of nuclei without dividing (**Figure 1**) (Petrucco et al., 2025). The propensity to bud or not to bud is influenced by cell density and nutrient availability (Goshima, 2022), but in a given environment cells can switch between budding and nuclear cycles stochastically. As the number of nuclei inherited by each bud is somewhat variable, mother cells can be found that have every integer number of nuclei from 1 to ∼20. In the same population, these proliferating cells span an impressive ∼100-fold size range, providing a tractable system to address whether changing cell size imposes different requirements for organellar activities.

**Figure 1:**
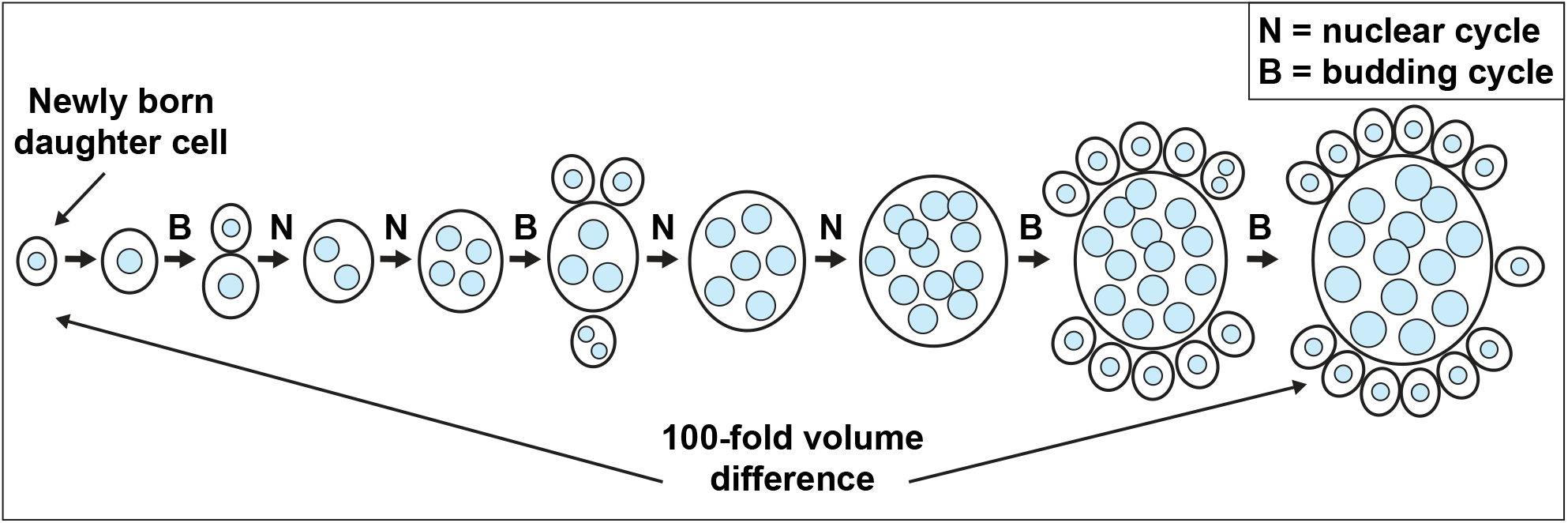
Schematic of *A. pullulans* budding proliferation. *A. pullulans* cells can undergo budding cycles (B) in which daughters usually receive 1 or 2 nuclei. Cells with more nuclei generally produce more buds. In addition, cells can undergo nuclear cycles (N) in which cells grow isotropically and mitosis doubles the nuclear content. Cells switch back and forth between budding cycles and nuclear cycles, resulting in cells of heterogeneous size and number of nuclei.

For the model budding yeast *Saccharomyces cerevisiae*, as for mammalian cells, the single nucleus limits the degree to which cells can enlarge while remaining healthy (Neurohr et al., 2019). In *A. pullulans*, nuclear number increases with size (Wirshing et al., 2024), presumably explaining how cells can remain healthy even when very large. However, the correlation between cell size and nuclear number was noisy, so that cells of similar size differed up to 4-fold in nuclear number, and conversely cells with the same number of nuclei differed up to 4-fold in volume. Interestingly, the total nuclear volume (sum of the volumes of every nucleus) exhibited a much tighter correlation with cell size, such that cells with higher nuclear density had smaller nuclei, and vice versa (Wirshing et al., 2024). This is consistent with current models positing that the abundance of nuclear macromolecules scales with cell volume, and determines the nuclear volume (Finan et al., 2009; Lemière et al., 2022; Rollin et al., 2023).

In a companion paper (Wirshing et al., 2026), we used fluorescent probes for mitochondria, ER, vacuoles, and peroxisomes in *A. pullulans* to measure how evenly these organelles were inherited by sibling buds emerging from the same mother. Here we characterize the scaling of these organelles with mother cell size. For mitochondria and ER, organelle density (organelle content per cell volume) remained constant in cells of very different sizes, while that was not the case for vacuoles and peroxisomes. For vacuoles, larger cells developed much larger central vacuoles that took up a larger fraction of the cell volume. For peroxisomes, larger cells developed fewer peroxisomes as a fraction of the cell volume, on average. Thus, in *A. pullulans* it appears that organelle composition adjusts to cell size.

## RESULTS

### Measuring organelle content and cell volume in *A. pullulans*

We used cytosolic 3xmCherry to enable 3D cell segmentation for volume measurement, and introduced organelle markers validated in the companion paper to quantify organelle content (Wirshing et al., 2024) (Wirshing et al., 2026). The markers are Cit1-GFP (mitochondria), Sec61-GFP and Elo3-GFP (ER), Vph1-GFP and Atg42-3xmNG (vacuole membrane and lumen, respectively), and Pex3-GFP (peroxisomes). Unless otherwise indicated, we report total fluorescence signal (GFP or mNG) as a measurement of total organelle content in the rest of the manuscript, and organelle density refers to the total organelle content divided by the cell volume.

### Mitochondrial content increases in parallel with cell volume

In *A. pullulans*, the mitochondrial network consists of interconnected tubules that extend throughout the cell volume (Wirshing et al., 2026, 2024), and this network appeared similar in cells of different sizes (**Figure 2A**). The total amount of mitochondria was proportional to cell volume (slope = 1 on a log-log plot) (**Figure 2B**), and the density of mitochondria was constant across the whole range of cell volumes (**Figure 2C and 2D**). In cells undergoing nuclear cycles without budding, volume growth was exponential with a doubling time of 169 ± 21 min (mean ± sd, n = 21 cells). Mitochondrial abundance increased continuously throughout the cell cycle (**Figure 2E and 2F, and Supplemental video 1**), yielding a consistent mitochondrial density with fluctuations not obviously linked to a particular cell cycle phase. Under these low-density growth conditions, cells switch between nuclear cycles (**Figure 2F**, red arrows mark mitoses) and budding cycles (**Figure 2F**, gray zones mark bud growth). During budding cycles, the mother cell volume remained constant, and the mitochondrial density decreased as mitochondria were transferred to buds (**Figure 2E**). These findings suggest that mitochondrial synthesis proceeds at a constant rate throughout the cell cycle that matches the cell’s growth rate. This does not appear to change as cells get bigger, so mitochondrial density is similar in small and large cells.

**Figure 2:**
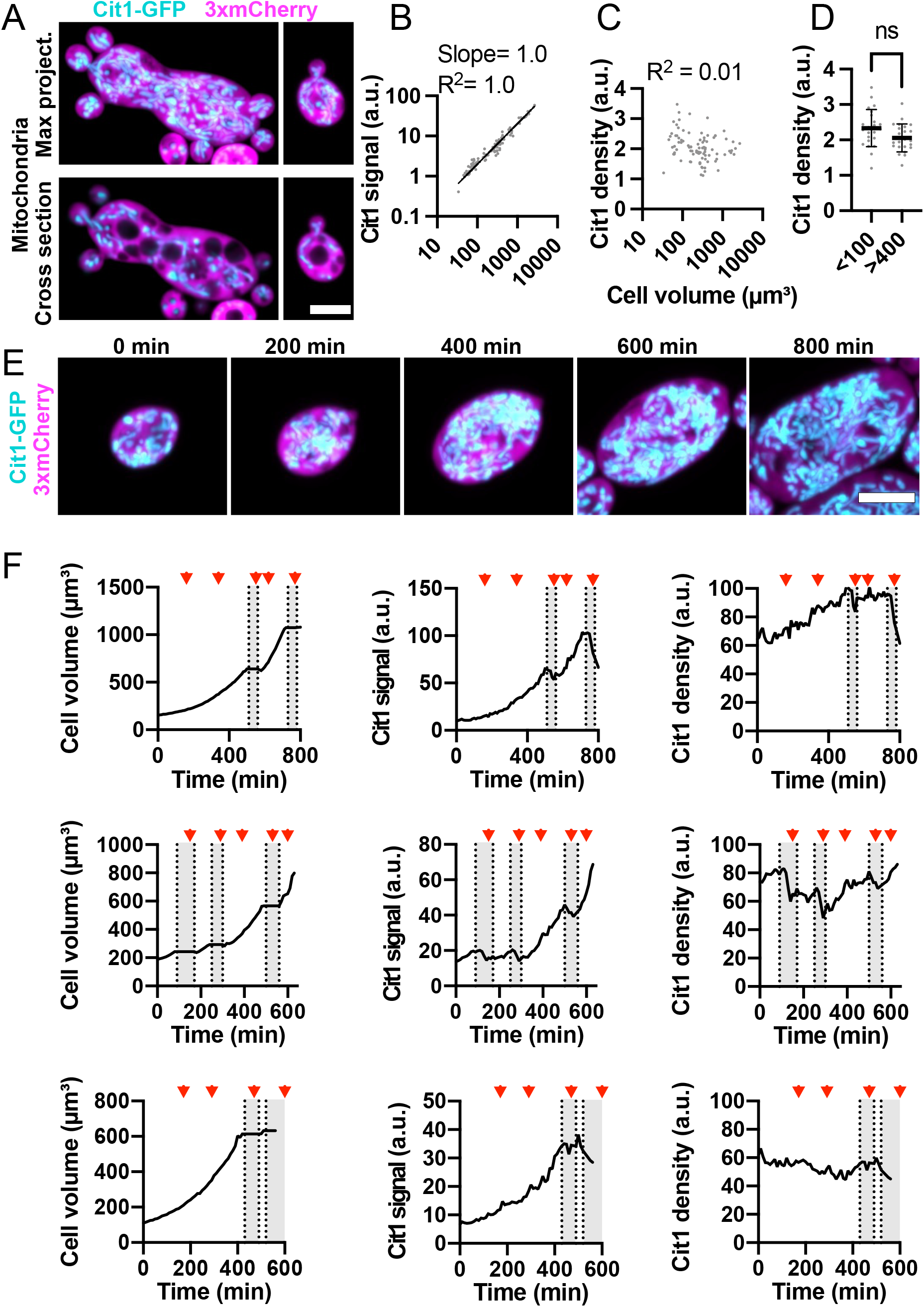
Mitochondria content scales linearly with cell volume. (**A**) Maximum intensity projection of a confocal Z-series (top) and single medial plane image (bottom) of large and small cells expressing a mitochondrial marker (cyan, Cit1-GFP) and cytosol marker (3xmCherry, magenta) (DLY25312). Scale bar, 5 µm. (**B**) Log-log plot of total mitochondrial content (Cit1-GFP signal) and cell volume. The best fit line, slope, and R^2^ values are shown (n = 91 cells). (**C**) Mitochondrial density (Cit1-GFP signal divided by cell volume) in the same cells analyzed in B. (**D**) Mitochondrial density for small (<100 µm^3^) and large cells (>400 µm^3^). Mean and standard deviation are indicated. p = 0.052 by student’s t-test (n = 23 small and 23 large cells). (**E**) Maximum intensity projection of a confocal time series of a cell from the same strain. Scale bar, 5 µm. (**F**) Quantification of cell volume (left), total mitochondrial content (middle), and mitochondrial density (right) for three representative cells. The top row is measured from the same cell shown in D. The volume measurements are only for the mother cell and do not count buds. Red arrowheads indicate the times when mitosis took place. Grey boxes indicate budded intervals.

### Total ER content increases in parallel with cell volume

The ER formed an extensive network connecting the nuclear envelope through thin tubules to sheets of ER at the cell cortex (cortical ER) (**Figure 3A and 3B**). Sec61-GFP decorated the tubules, sheets, and nuclear envelopes in a homogeneous manner, while Elo3-GFP was more concentrated at the nuclear envelope (**Figure 3B**). Large cells expressing Sec61-GFP also contained bright globular structures that were not observed in smaller cells (**Figure 3A arrows**). These structures were less prevalent in cells expressing Sec61-mCherry, and were not visible in cells expressing Elo3-GFP (compare **Figure 3A and 3B**). Thus, it is possible that the structures were in some way induced as an artifact of tagging Sec61. Alternatively, they might be physiological structures that tend to exclude tagged Elo3 (and to a lesser extent mCherry-tagged Sec61). At present, we cannot definitively distinguish between these possibilities, so we proceeded to quantify ER abundance using both probes. ER quantification using Sec61-GFP (**Figure 3 and 4**) or Elo3-GFP (**Supplemental Figure 1**) yielded similar conclusions.

**Figure 3:**
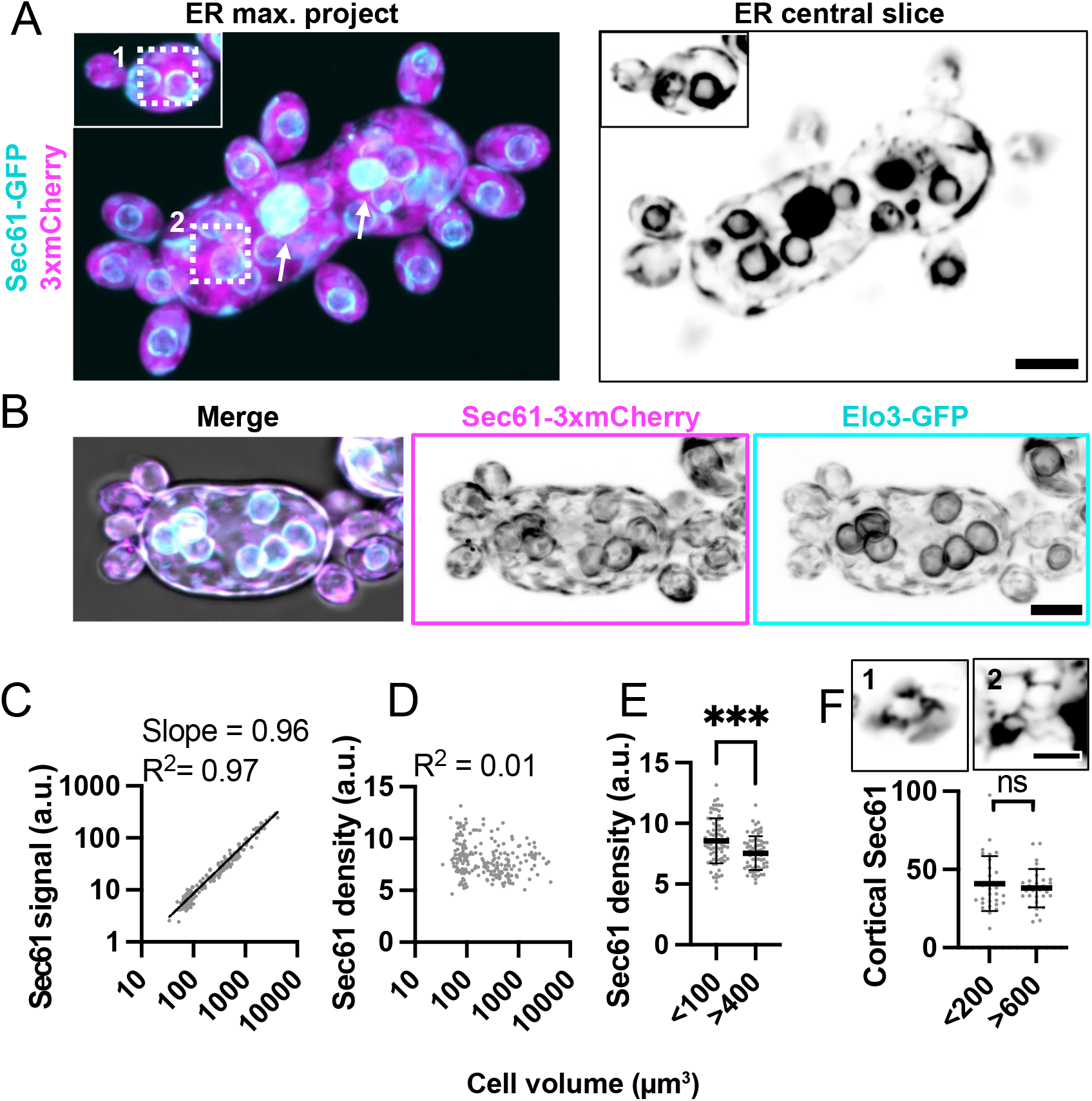
ER content scales linearly with cell volume. (**A**) Maximum intensity projection of a confocal Z-series (top) and single medial plane image (bottom) of large and small cells expressing an ER marker (cyan, Sec61-GFP) and cytosol marker (3xmCherry, magenta) (DLY24944). The single slice is shown in greyscale to highlight fine ER structures. Arrows indicate large globular structures. Dashed boxes indicate inserts shown in E. Scale bar, 5 µm. (**B**) Maximum intensity projection of a confocal Z-series of a cell expressing two different ER markers, Sec61-3xmCherry (magenta) and Elo3-GFP (cyan) (DLY27440). Grayscale images illustrate higher Elo3 concentration at the nuclear envelope. Scale bar, 5 µm (**C**) Log-log plot of total ER content (Sec61-GFP signal) and cell volume. The best fit line, slope, and R^2^ values are shown (n = 209 cells). (**D**) ER density (Sec61-GFP signal divided by cell volume) in the same cells analyzed in C. (**E**) ER density for small (<100 µm^3^) and large cells (>400 µm^3^). Mean and standard deviation are shown. p ≤ 0.001 by student’s t-test (n = 78 small and 65 large cells). (**F**) Mean cortical ER intensity measured from glancing slices in small (<200 µm^3^) and large (>600 µm^3^) cells. The mean and standard deviation are shown. p = 0.464 by student’s t-test (n = 30 small and 30 large cells). Insets show a single confocal glancing slice of the cortical ER from the regions in the dashed boxes in A. Scale bar, 2 µm.

**Figure 4:**
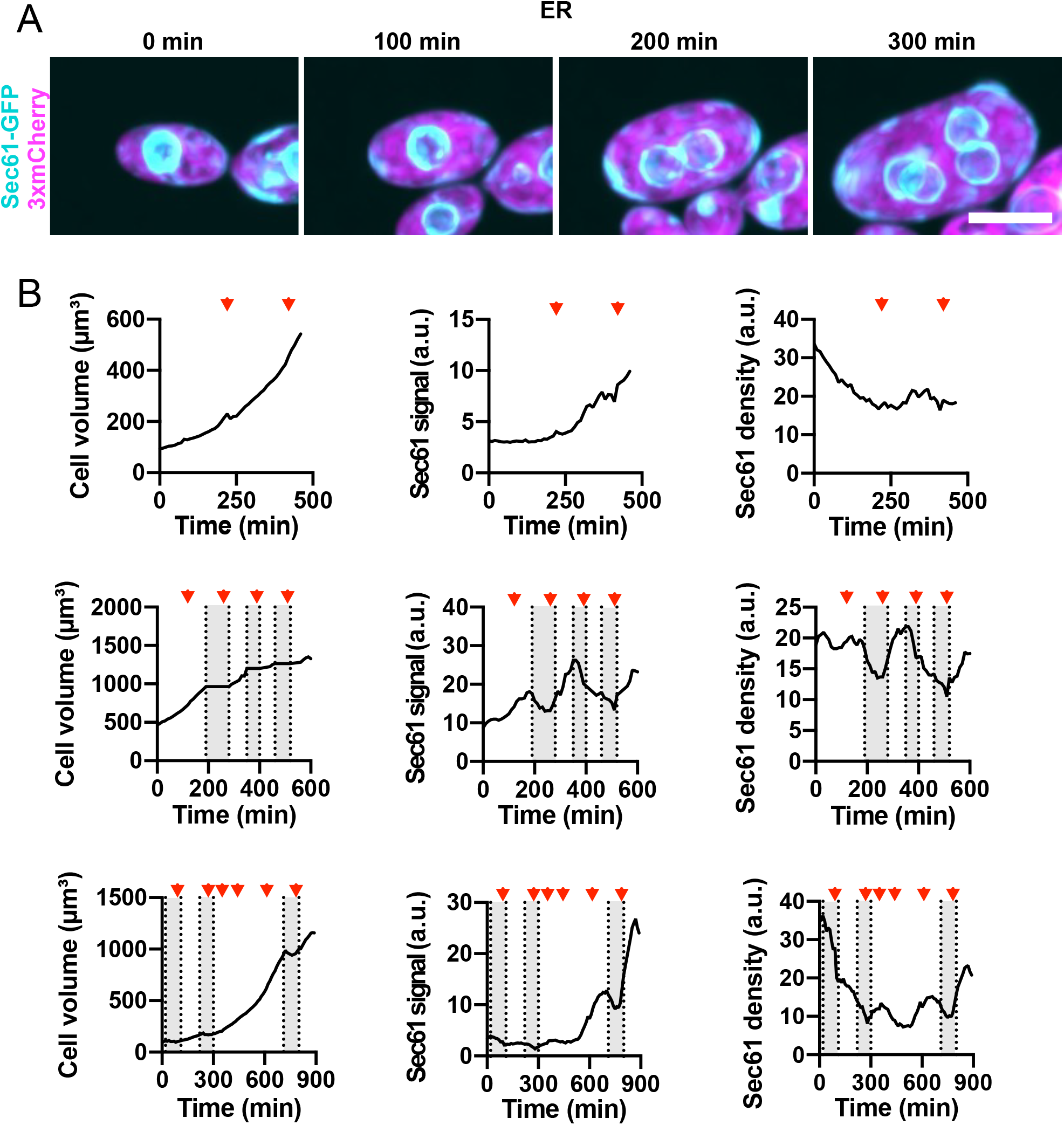
ER content increases as cells grow. (**A**) Maximum intensity projection of a confocal time series of a cell expressing ER marker Sec61-GFP (cyan) and cytosol marker 3xmCherry (magenta) (DLY24944). Scale bar, 5 µm. (**B**) Quantification of cell volume (left), total ER content (middle), and ER density (right) for three representative cells. The top row is measured from the same cell shown in A. Red arrowheads indicate the times when mitosis took place. Grey boxes indicate budded intervals.

As with mitochondria, ER content scaled linearly with cell volume (slope = 0.96 on a log-log plot) (**Figure 3C**). ER density remained fairly constant across a range of cell volumes and there was only a slight difference in density between small (<100 µm3) and large (>400 µm3) cells (**Figure 3D and 3E**). The average ER density measured with Sec61-GFP was slightly lower in large cells (**Figure 3D**), but this was not the case for Elo3-GFP (**Supplemental Figure 1B**). Because the surface area to volume ratio (SA/V) decreases with increasing volume in ovoid cells, we wondered whether cortical ER would scale with cell volume or surface area. Measuring Sec61-GFP in single glancing cortical slices suggested that cortical ER density remains similar in cells of different sizes (**Figure 3F**). This suggests that the amount of cortical ER scales with the cell’s surface area, while total amount of ER scales with the cell’s volume. Thus, the fraction of ER present in cortical sheets would have to decrease as cells grow.

Monitoring ER content as cells grew revealed larger fluctuations in ER density than what we had observed for mitochondria (**Figure 4 and Supplemental video 2**). In some cases, cells grew without increasing ER content as measured using Sec61-GFP and then accumulated ER more rapidly (**Figure 4B**). This is at odds with our finding that ER content scales with cell volume for cells grown in liquid media (**Figure 3C**). We speculate that transfer to agarose slabs for long-term imaging may induce stress in some of the cells that then transiently delays ER accumulation. While this introduces some noise, the overall pattern of ER accumulation in most cells was continuous throughout the cell cycle, as for mitochondria (**Figure 4B, Supplemental Figure 1B**).

### Vacuole content increases more than cell volume as cells grow larger

All cells had multiple vacuoles of varying sizes, but the biggest vacuoles were generally larger in larger cells (**Figure 5A**). Total vacuole membrane (Vph1-GFP signal) scaled with cell volume (slope = 1.1 on a log-log plot) (**Figure 5B**), and the vacuole membrane density was similar in cells of different sizes (**Figure 5C and 5D**). Monitoring vacuole membrane as cells grew confirmed that vacuole membrane increased in parallel with cell volume (**Figure 5E and 5F and Supplemental video 3**). During budding cycles, vacuole content dipped in mother cells as vacuoles were transferred to buds, but during nuclear cycles vacuole membrane content increased continuously (**Figure 5E and 5F**).

**Figure 5:**
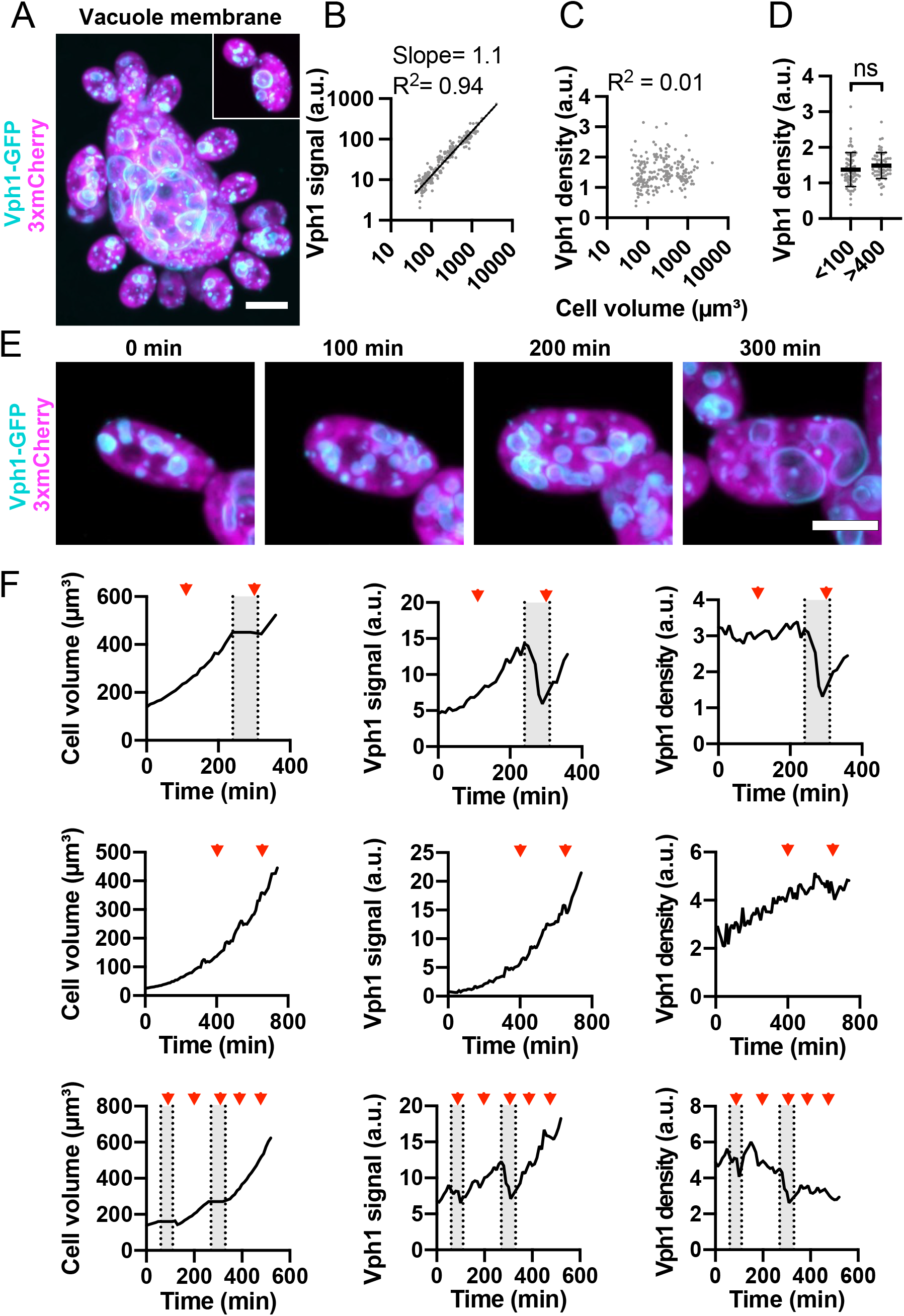
Vacuole membrane content scales linearly with cell volume. (**A**) Maximum intensity projection of a confocal Z-series showing differently sized cells expressing the vacuole membrane marker Vph1-GFP (cyan) and the cytosol marker 3xmCherry (magenta) (DLY24947). Scale bar, 5 µm. (**B**) Log-log plot of total vacuole membrane content (Vph1-GFP signal) and cell volume. The best fit line, slope, and R^2^ values are shown (n = 217 cells). (**C**) Vacuole membrane density (Vph1-GFP signal divided by cell volume) in the same cells analyzed in B. (**D**) Vacuole membrane density in small (<100 µm^3^) and large cells (>400 µm^3^). Mean and standard deviation are shown. p = 0.128 by student’s t-test (n = 71 small and 61 large cells). (**E**) Maximum intensity projection of a confocal time series of a cell expressing vacuole membrane marker Vph1-GFP (cyan) and cytosol marker 3xmCherry (magenta) (DLY24947). Scale bar, 5 µm. (**F**) Quantification of cell volume (left), total vacuole membrane content (middle), and vacuole density (right) for three representative cells. The top row is measured from the same cell shown in E. Red arrowheads indicate the times when mitosis took place. Grey boxes indicate the budded intervals.

We next used the vacuole lumen marker, Atg42-3xmNG, to quantify the relationship between cell and vacuole volumes. We found there was not a straightforward power law scaling relation between total Atg42-3xmNG signal and cell volume (**Figure 6A**). The density of Atg42-3xmNG increased with cell volume (**Figure 6B**), suggesting that total vacuole volume grew faster than cell volume. Using the Atg42-3xmNG signal to measure vacuole volumes (see methods for details) confirmed this conclusion (**Figure 6C and 6D**), and in both cases small cells (<100 µm3) had significantly lower vacuole density than large ones (>400 µm3) (**Figure 6E and 6F**). Looking at individual vacuoles in differently sized cells, it was apparent that in large cells, a few central vacuoles became much larger than the rest (>10-fold) (**Figure 6G**). Thus, as cells increase in size, the number of vacuoles increases, and a few central vacuoles become much larger in volume resulting in vacuoles occupying a higher fraction of the total cell volume.

**Figure 6:**
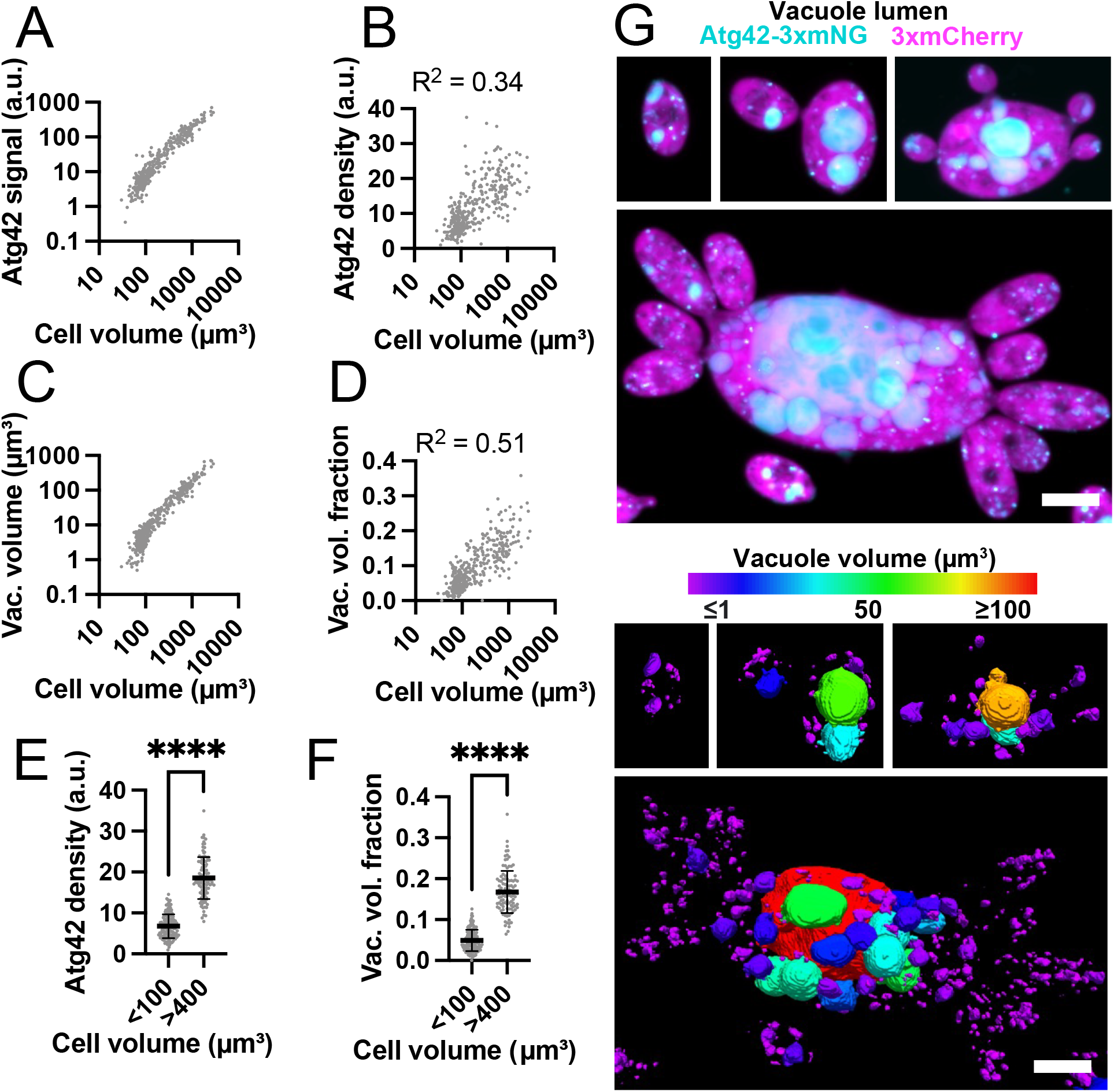
Vacuoles occupy a greater fraction of the cell volume in large cells. (**A**) Log-log plot of total vacuole lumen content (Atg42-3xmNG signal) and cell volume (n = 375 cells) (DLY25963). (**B**) Vacuole lumen density (Atg42-3xmNG signal divided by cell volume) in same cells analyzed in A. An F test reveals the slope of the best fit line (not shown) is significantly different from zero (p = <0.0001). (**C**) Log-log plot of total vacuole volume (calculated based on segmentation of Atg42-3xmNG signal) and cell volume for the same cels analyzed in A. (**D**) Vacuole volume fraction (total vacuole volume divided by cell volume) for the same cells analyzed in A. (**E**) Vacuole density (Atg42-3xmNG signal divided by cell volume) for small (<100 µm^3^) and large cells (>400 µm^3^). Mean and standard deviation are shown. ****, p ≤ 0.0001 by student’s t-test (n = 144 small and 104 large cells). (**F**) Vacuole volume fraction for the same cells analyzed in E. ****, p≤ 0.0001 by student’s t-test. (**G**) Top: Maximum intensity projection of a confocal Z-series of variously sized cells expressing the vacuole lumen marker Atg42-3xmNG (cyan) and cytosol marker 3xmCherry (magenta) (DLY25963). Scale bar, 5 µm. Bottom: The same cells shown with the vacuoles false colored according to vacuole volume.

### Peroxisome density and variability is higher in small cells

Unlike vacuoles, individual peroxisomes were of similar size in small and large cells (**Figure 7A**). In contrast to the other organelles, total peroxisome content (Pex3-GFP signal) subscaled with cell volume (slope 0.75 on a log-log plot) (**Figure 7B**), so that peroxisome density decreased with increasing cell volume (**Figure 7C**). Cell-to-cell variability in peroxisome density was also lower in large cells (>400 µm3) than small ones (<100 µm3) (**Figure 7D**).

**Figure 7:**
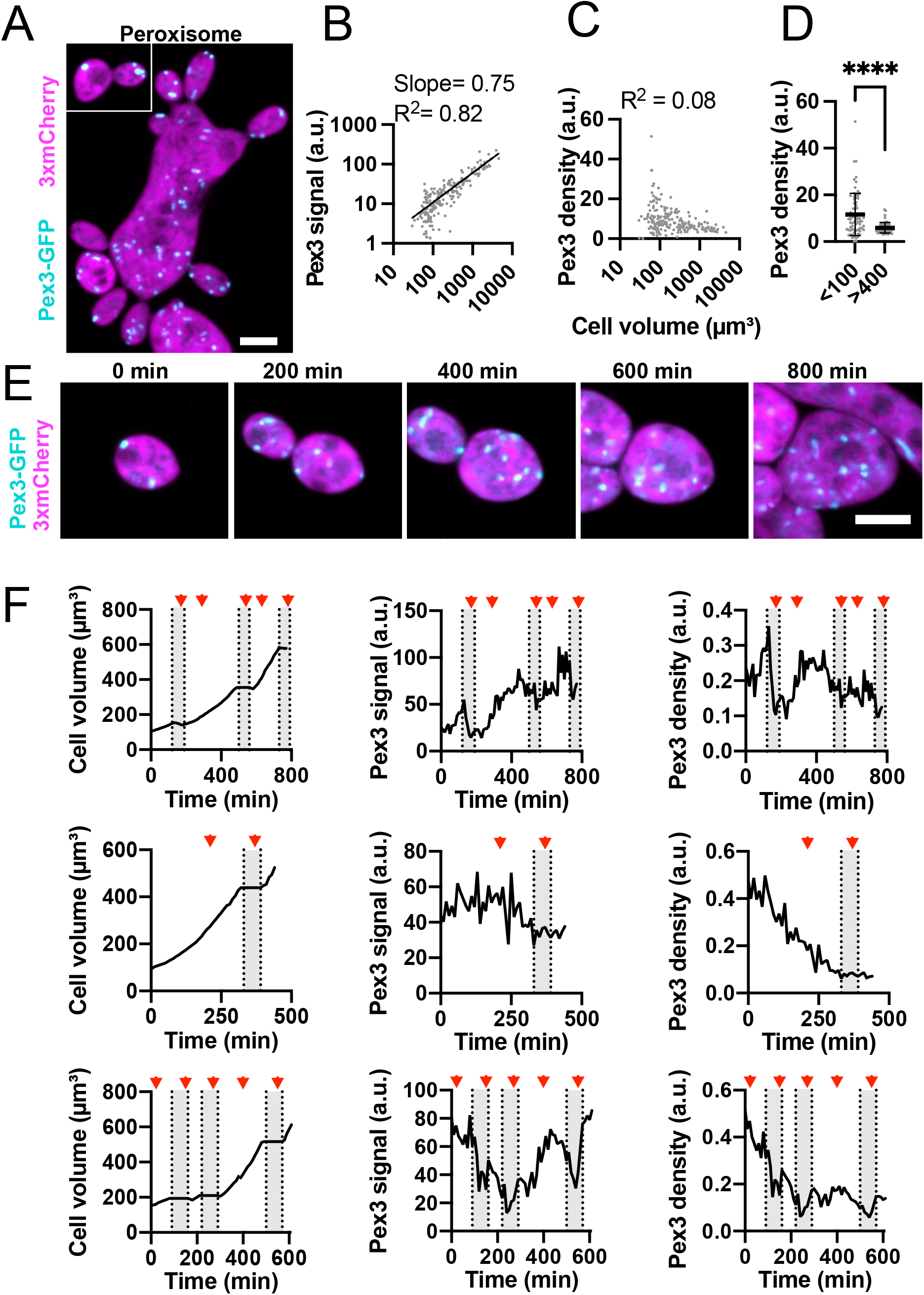
Peroxisome content heterogeneity decreases with increasing cell size. (**A**) Maximum intensity projection of a confocal Z-series of differently sized cells expressing a peroxisome marker (Pex3-GFP, cyan) and cytosol marker (3xmCherry, magenta). Scale bar, 5 µm (DLY25001). (**B**) Log-log plot of total peroxisome content (Pex3-GFP signal) and cell volume. The best fit line, slope, and R^2^ values are shown (n = 206 cells). (**C**) Peroxisome density (Pex3-GFP signal divided by cell volume) in the same cells analyzed in B. (**D**) Peroxisome density in small (<100 µm^3^) and large (>400 µm^3^) cells (n = 73 small and 55 large cells). ****, p ≤ 0.0001 by student’s t-test with Welch’s correction for unequal variance. The variance in large cells is significantly smaller (p < 0.0001 by F-test). (**E**) Maximum intensity projection of a confocal time series of a cell expressing Pex3-GFP (cyan) and 3xmCherry (magenta). Scale bar, 5 µm. (**E**) Quantification of cell volume (left), total peroxisome content (middle), and peroxisome density (right) for three representative cells. The top row is measured from the same cell shown in D. Red arrowheads indicate the times when mitosis took place. Grey boxes indicate the budded intervals.

Peroxisome content in individual cells showed variable patterns. In some cases, peroxisome content and cell size increased in parallel, maintaining a constant peroxisome density (**Figure 7E and 7F, top row**). In other cases, peroxisome content did not change as the cell grew, so that peroxisome density decreased with time (**Figure 7F, bottom two rows**). In budding cycles, peroxisome content in mother cells decreased as peroxisomes were transferred to buds (**Figure 7F**). In all cases, peroxisome accumulation appeared unrelated to cell cycle phase (**Figure 7F**). Thus, peroxisome density can fluctuate considerably, and tends to decrease as cells grow.

## DISCUSSION

A considerable body of evidence supports the idea that among cells in a proliferating population, organelle content scales with cell volume (Banerjee and Banerjee, 2025; Marshall, 2020; Wesley et al., 2020). This makes intuitive sense, as larger cells would make more secretory proteins (necessitating more ER), use more energy (necessitating more mitochondria), and so on. However, given the small size of many organelles, it is difficult to measure organelle contents via light microscopy with great precision. Thus, small deviations from proportional scaling are easy to miss, particularly for typical cells that populate a narrow size range. The unusual proliferation strategy of *A. pullulans*, with both nuclear cycles and budding cycles, generates cells that vary over a much wider size range (**Figure 1**). This provides an opportunity to detect departures from linear scaling, and to ask whether the inherent differences in biophysical properties of large and small cells might lead to different organelle requirements. Here, we characterize the scaling of four organelles (mitochondria, ER, vacuole, and peroxisomes) with cell size in *A. pullulans*, and report that different organelles exhibit different behaviors.

### Mitochondria

In *A. pullulans* the mitochondrial network was dispersed throughout the cell volume and appeared similar in both small and large cells. This is distinct from *S. cerevisiae* where mitochondria are excluded from the cell center and localize to the cell periphery (Lazzarino et al., 1994; Perktold et al., 2007), but similar to several other ascomycete fungi where mitochondria extend throughout the cell (Faoro et al., 2022; Fekete et al., 2007; Mims et al., 1988; Riquelme et al., 2014). One potential advantage to having a distributed mitochondrial organization is that it allows for ATP production throughout the cell volume. For small *S. cerevisiae* cells (∼5 µm diameter) where ATP needs to diffuse short distances, the ATP concentration is fairly unform (Takaine et al., 2019). However, for larger cells the ATP concentration can vary and be higher where mitochondria are clustered (Kumar and Johnston, 2025; Schuler et al., 2017). Therefore, the distributed mitochondrial organization in *A. pullulans* potentially ensures adequate ATP availability in cells of various sizes and shapes.

Mitochondrial content increased proportionally with cell volume, maintaining a constant density of mitochondria in the cell. Similar results have been reported for *S. cerevisiae* and animal cells (Miettinen and Björklund, 2016; Posakony et al., 1977; Rafelski et al., 2012). Interestingly, in several animal cells mitochondria functionality decreases as cells increase in volume (Miettinen and Björklund, 2016), so that cells above a certain size optimum are less fit. It will be interesting to test whether and how *A. pullulans* circumvents this issue.

### ER

The ER in *A. pullulans* formed an extensive network linking the nuclear envelope(s) to cortical sheets at the plasma membrane. Similar to the mitochondria, total ER content increased in proportion to cell volume. However, the cortical ER content increased in proportion to cell surface area, such that fraction of the plasma membrane associated with ER sheets remained constant. Thus, in larger cells a smaller fraction of the total ER was present in cortical sheets. Cortical ER is tethered to the plasma membrane at contact sites that regulate lipid and Ca2+ homeostasis (Li et al., 2021; Wu et al., 2018; Zaman et al., 2020). Such contact sites are formed by multiple protein complexes that dynamically regulate the density of ER-PM contacts to meet cell needs (Li et al., 2021;

Wu et al., 2018; Zaman et al., 2020). How ER-PM contact sites are regulated in *A. pullulans* so as to maintain a constant density of ER at the plasma membrane across a large range of cell size remains an intriguing open question.

### Vacuoles

Vacuoles in *A. pullulans* were globular and variable in size. Notably, larger cells contained a subset of vacuoles that grew to a much larger size than the rest, such that the largest vacuoles in big cells were larger than the entire volume of many smaller cells. Thus, vacuole size is to some extent dependent on cell size. Total vacuolar membrane scaled with cell volume, and vacuoles in larger cells accounted for a larger fraction of the total cell volume. Similarly, in *S. cerevisae* vacuole surface area density remains constant while the vacuole volume fraction increases with increasing cell volume (Chan and Marshall, 2014).

If cells had a fixed number of spherical vacuoles that grew larger as cells became larger, then scaling of vacuole membrane with cell volume (slope =1 on log-log plot) would be accompanied by superscaling of vacuole volume with cell volume (slope =1.5 on log-log plot). Conversely, if cells kept vacuole volumes constant and simply increased the number of vacuoles with cell volume, then both vacuole membrane and vacuole volume would scale linearly with cell volume. Neither of these scenarios applies to *A. pullulans*. Both vacuole numbers and vacuole volumes increased with cell volume in this system. Although the fraction of total cell volume occupied by vacuoles increased continuously with cell size (from ∼2% in small cells to ∼20% in large cells), the increase did not follow a simple power law scaling. Instead, the vacuole volume fraction grew more rapidly in smaller cells than in larger cells. This may be related to the heterogeneous distribution of vacuole sizes within cells.

It is not immediately apparent why some vacuoles grow much larger. In *S. cerevisiae*, vacuoles exhibit both fusion and fission (Weisman, 2003), so it may be that the balance between these processes changes as cells become larger. However, the fact that vacuole sizes within a cell can be very heterogeneous suggests that there is some autonomous control of the process by the vacuole itself, rather than a target size set by the cell. It is also unclear why total (summed) vacuole volume would increase as a fraction of cell volume in larger cells. One possibility is that expanding the vacuole enables these cells to mitigate the changing cytoplasm-to-surface area ratio as cells grow larger (Okie, 2013). Moreover, vacuole size may impact the multiple functions of the organelle, including protein/organelle degradation and ion homeostasis (Li and Kane, 2009).

### Peroxisomes

Peroxisomes in *A. pullulans* were small and globular, and larger cells did not contain larger peroxisomes (just more peroxisomes of similar size). Peroxisome content was extremely variable in small cells, but became more homogenous in larger cells. The high peroxisome content variability in small cells likely reflects the variability in peroxisome inheritance by sibling daughters (Wirshing et al., 2026). If peroxisomes proliferated by exponential division of pre-existing peroxisomes, then we would expect the starting variability to be maintained or even increased as cells grew larger. Instead, daughters inheriting much more (or fewer) peroxisomes than average seemed to adjust peroxisome content to become more consistent as they grew. This may simply reflect the statistical principle that variability in number decreases for objects present at higher mean numbers. We note that peroxisome content fluctuated much more markedly than other organelles even when considering a single cell growing over time. Thus, it does not appear that peroxisomes are under tight control in our growth conditions.

Unlike the other organelles, peroxisome content increased sublinearly with cell volume. While the function of this is unknown, peroxisomes have multiple roles including fatty acid β-oxidation and regulation of reactive oxygen species homeostasis (Sibirny, 2016). In multiple fungal systems peroxisome content is dramatically increased or decreased depending on the environment (Goodman et al., 1990; Gurvitz and Rottensteiner, 2006; Tuttle et al., 1993; Veenhuis et al., 1987, 1978). As in other fungi, *A. pullulans* cells may dynamically adjust peroxisome content to an ideal setpoint for a given environment. The ability to detect and adjust peroxisome content may explain why it is unnecessary for mother cells to apportion the same amount of peroxisomes to each daughter cell.

### Conclusions

The importance of having the right organelle content, and the mechanisms by which cells regulate organelle contents, are only beginning to be examined (Marshall, 2020). Proliferating *A. pullulans* cells populate an impressive range of sizes, allowing an investigation into whether cells of different sizes might exhibit preferences for particular organelle contents. We found that the relationship between cell volume and organelle content was different for different organelles. The density and organization of the mitochondria and ER remained fairly constant as cells grew larger, while vacuoles took up an increasing fraction, and peroxisomes took up a decreasing fraction, of the total cell volume in larger cells. This raises the question of how and why *A. pullulans* cells adjust organelle content as they grow.

## METHODS

### *A. pullulans* strains and maintenance

All strains used in this study are in the *Aureobasidium pullulans* EXF-150 strain background (Gostinčar et al., 2014). A list of all strains with genetic modifications is provided in **Supplemental Table 1**. *A. pullulans* was grown at 24°C in standard YPD medium (2% glucose, 2% peptone, 1% yeast extract) with 2% BD Bacto^™^ agar (214050, VWR) in plates, unless otherwise indicated.

### Strain construction

Construction of strains expressing the cytoplasmic marker (mCherry) and organelle markers Cit1-GFP, Sec61-GFP, Vph1-GFP, Atg42-3xmNG, and Pex3-GFP were described previously (Wirshing et al., 2026). All organelle markers are C-terminally tagged at their native loci. To determine compare the two ER markers Sec61 and Elo3, a Sec61-3xmCherry marker was first introduced at the native Sec61 locus and then an Elo3-GFP marker was added to this background. Integration was targeted to the loci of interest by 3-part-PCR using pAPInt vectors with GFP or three tandem copies of mCherry and hygromycin or nourseothricin drug resistance cassettes as described previously (Colarusso et al., 2025). Briefly, 1 kb homology regions flanking the desired insertion site were amplified from the genome along with the pAPInt integration cassette (fluorescent probe + selection cassette) with 60-70 bp overlap between each PCR product. PCR products were pooled, cleaned, and concentrated to ∼1,500 ng/µl as described previously (Wirshing et al., 2024). Cleaned PCR products were added directly to *A. pullulans* competent cells in equimolar ratios, and cells were transformed using PEG/LiOAc/ssDNA as described previously (Wirshing et al., 2024). Transformants were selected on YPD plates supplemented with Hygromycin B (400051-1MJ, Millipore) at a final concentration of 174 µl/l (∼70.4 mg/l) or 50 mg/l Nourseothricin (N1200-1.0, Research Products International). Colonies appearing after 2-3 days at 24°C were screened for the desired genome modification by colony PCR using the Phire Plant Direct PCR Master Mix (#F-160S, Thermo Scientific) following the manufacturer’s instructions. Positive colonies were also confirmed to show the expected fluorescent signals.

### Live-cell imaging and image analysis

For single-timepoint snapshots used to compare organelle content and cell size across a proliferating population, a single colony was used to inoculate 5 ml of YPD (2% glucose) and the culture was grown overnight at 24°C to a density of 1-5 × 106 cells/ml. At these intermediate cell densities, *A. pullulans* cells undergo a mixture of nuclear cycles and budding cycles, so that cells exhibit a mixture of different sizes. At higher cell densities, budding cycles dominate and most cells are small, while at lower cell densities, more hyphae are present. Overnight cultures were concentrated to a density of ∼7 × 107 cells/ml by briefly spinning at 9391 rcf for 10 s and removing the supernatant. 2 µl of the concentrated cells were added to the bottom of a glass-bottomed 8-well Ibidi chamber (80827, Ibidi) and covered with a 200 mg 5% agarose (97062-250, VWR) pad made with CSM (6.71 g/L BD DifcoTM Yeast Nitrogen Base without Amino Acids, BD291940, FisherScientific, 0.79 g/L Complete Supplement Mixture, 1001-010, Sunrise Science Products, and 2% glucose). These cells grown in liquid culture were then imaged immediately (see settings below) to capture still images of variously sized cells expressing the different organelle markers.

For long-term live-cell imaging, overnight cultures were harvested at slightly lower densities (1-3 × 106 cells/ml). Importantly, cells were also mounted in the Ibidi chambers at a lower density. This is key to promote nuclear cycles. The cell density of the overnight culture was adjusted to 2 × 106 cells/ml, and 2 µl of cells were mounted in an Ibidi chamber as described above and similarly covered with an agarose pad made with 7% agarose instead of 5%. Time-lapse microscopy (see settings below) was then used to monitor organelle content as cells grew. All imaging was conducted at room temperature (20-22°C).

Cells were imaged on a Nikon Ti2E inverted microscope with a CSU-W1 spinning-disk head (Yokogawa), CFI60 Plan Apochromat Lambda D 60x Oil Immersion Objective (NA 1.42; Nikon Instruments), a LUN-F compact four-line laser source (405 nm 100 mW, 488 nm 100 mW, 561 nm 100 mW, and 640 nm 75 mW fiber output), and a Hamamatsu ORCA Quest qCMOS camera controlled by NIS-Elements software (Nikon Instruments). For still images, the entire cell volume was acquired using 79-89 Z-slices at 0.2 μm step intervals. Exposure times of 50 ms at 50% laser power were used with excitation at 488 nm (with ET525/36m Emission Filter) to excite GFP or mNG, and with excitation at 561 nm (with ET605/52m Emission Filter) to excite mCherry. To track cell growth and organelle content over time, Z-stacks (17 slices, 0.7 µm interval) were acquired every 10 minutes for 12-16 hours using 50 ms exposures at 15% laser power for excitation 488 nm and 10% laser power for excitation at 561 nm.

To characterize the relationship between total cell volume and organelle content, confocal Z-stacks of each organelle marker strain were denoised and 3D segmented using NIS-Elements General Analysis 3 software (GA3, Nikon Instruments). The cytoplasmic mCherry signal was used to generate a mask of the cell and to measure cell volumes. The cell mask was also used to subtract the background signal by measuring the average GFP or mNG signal outside of the cell mask and subtracting this value from each image. Next, the organelle GFP or mNG signal was used to 3D segment organelle volumes to generate an organelle mask. The cytoplasmic GFP or mNG background signal in each cell was subtracted by setting the intensity value of any pixels within the cell but outside of the organelle mask to zero. To calculate the total organelle (GFP or mNG) signal in each cell, the GFP or mNG pixel values (after background subtraction) in each cell volume were summed. Here we report total GFP or mNG signal within each cell as total organelle content. The organelle density was calculated by dividing this total organelle content by the cell volume.

To measure the amount of cortical ER, images were processed and segmented as described above. The first Z-slice where the ER signal was in focus at the cell cortex was manually determined by scrolling through the Z-stack. To compare ER cortical densities in cells of different sizes, we initially used the same 100 µm^3^ cutoff for “small cells” as we used in the rest of the figures. However, for several cells smaller than 100 µm^3^, the area of the cortex in focus in a single slice was quite small. ER density at the cortex is variable (**Figure 3F**). To avoid error from sampling only a small area, we measured the cortical signal in cells <200 µm^3^. To enable comparison with cells at least 3-fold larger in volume, we compared these measurements to those from cells >600 µm^3^, instead of the >400 µm^3^ cutoff used for “large cells” in other figures. The mean Sec61-GFP or Elo3-GFP signal (after background subtraction) was measured from the single glancing slice using the mask of the 3xmCherry signal to mark the cell boundaries. We report the mean signal from this slice as a measurement of the density of ER at the cortex.

To measure vacuole volumes, images of cells expressing the luminal marker Atg42-3xmNG were 3D deconvolved in NIS-Elements General Analysis 3 software (GA3, Nikon Instruments) before 3D segmentation. The size of vacuole compartments in cells is highly variable with some compartments appearing to have diameters below the light microscopy resolution limit (∼200-250 nm). Because the actual diameters of these small compartments cannot be accurately measured by light microscopy, compartments with diameters <460 nm were excluded from our volume measurements. Thus, we potentially underestimate the true vacuole volume fraction in cells. Despite this, there is a strong correlation between total Atg42-3xmNG signal and vacuole volumes (R2 = 0.94) (**Supplemental Figure 2**).

To track isotropic growth and organelle accumulation captured in time lapse experiments, NIS-Elements General Analysis 3 (GA3, Nikon Instruments) software was used to track growing cells and cells were segmented and measured as described above for analysis of still images.

### Statistical analysis

All statistical analysis was done using GraphPad Prism. Unless indicated otherwise, data distributions were assumed to be normal, but this was not formally tested. Statistical comparison between indicated conditions was conducted using the two-sided Student’s t test as indicated in the figure legends. Where indicated, a two-sample F-test was used to test for significant differences between dataset variances. We used linear regression to test for correlations between organelle densities and cell volumes, and we used a regression F-test to determine if the slope was significantly different from zero; when the p-value was significant, this is indicated. In all cases, differences were considered significant if the p-value was <0.05.

## ABBREVIATIONS

ER: Endoplasmic reticulum
ER-PM: Endoplasmic reticulum-plasma membrane
GFP: Green fluorescent protein
PM: mNeonGreen: mNG Plasma membrane
SA/V: Surface area to volume ratio

## ACKNOWLEDGEMENTS

We thank Stephen P. Bell and members of the Lew Lab for comments on the manuscript. This work was funded by NIH/NIGMS grant R35GM122488 to D.J.L.

## AUTHOR CONTRIBUTIONS

Conceptualization, review, and editing manuscript— A.C.E. Wirshing, and D.J. Lew. Data curation, investigation, methodology, visualization, validation, and formal analysis—A.C.E. Wirshing. Drafting of manuscript— A.C.E. Wirshing. Project administration, supervision, and resources— D.J. Lew

## SUPPLEMENTAL FIGURE CAPTIONS

**Supplemental Figure 1:**
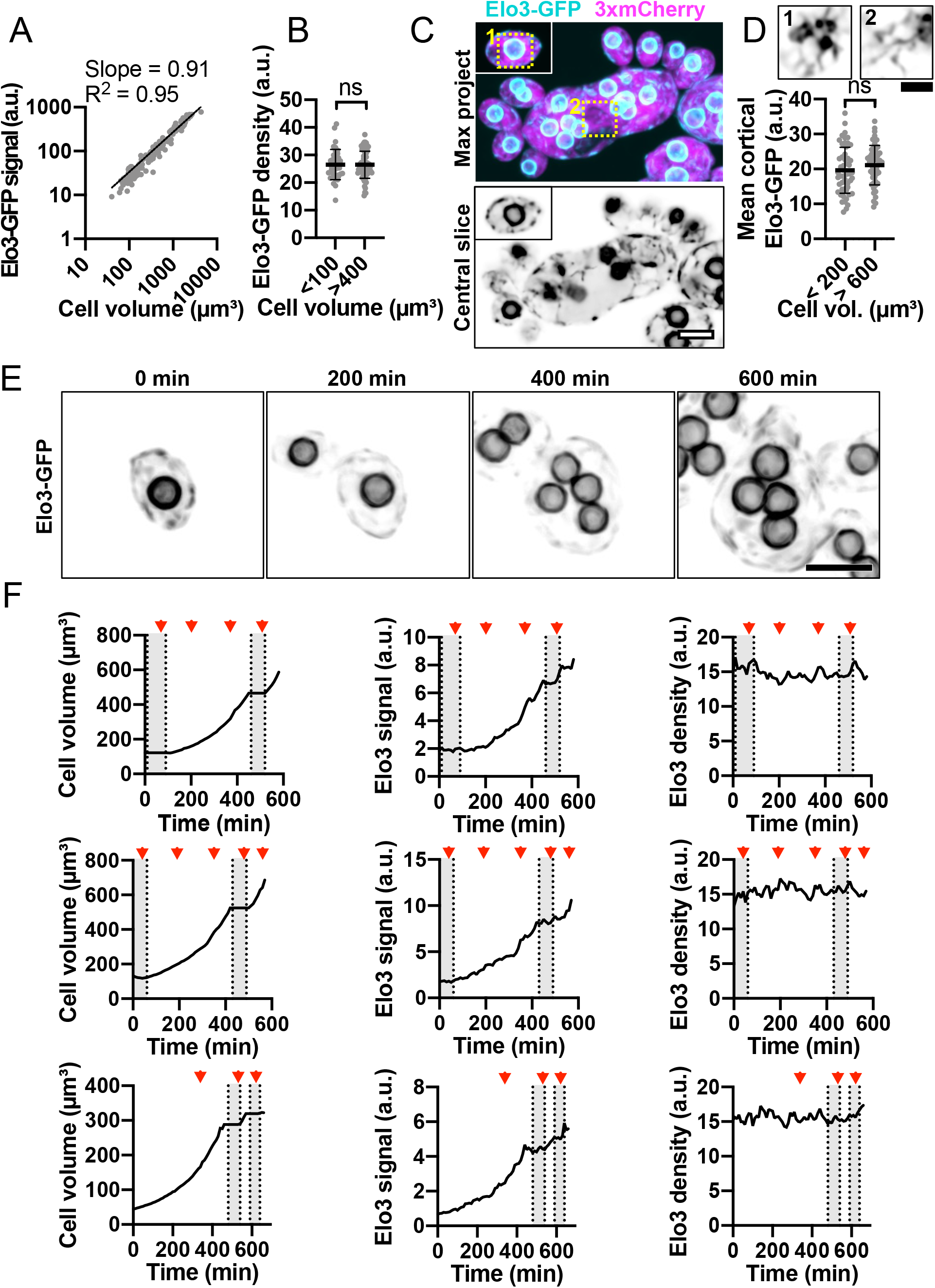
ER content scales linearly with cell volume when Elo3-GFP is used to visualize the ER. (**A**) Log-log plot of total ER content (Elo3-GFP signal) and cell volume. The best fit line, slope, and R^2^ values are shown (n = 179 cells). (**B**) ER density for small (<100 µm^3^) and large cells (>400 µm^3^). Mean and standard deviation are shown. p = 0.221 by student’s t-test (n = 38 small and 70 large cells). (**C**) Maximum intensity projection of a confocal Z-series (top) and single medial plane image (bottom) of cells of different sizes expressing the ER marker Elo3-GFP (cyan) and cytosol marker 3xmCherry (magenta) (DLY27372). The central slice is shown in greyscale to highlight fine ER structures. Dashed boxes indicate insets shown in D. Scale bar, 5 µm. (**D**) Top: Example confocal single glancing slice images of cortical ER from the regions in the dashed boxes in C. Bottom: mean cortical ER intensities measured from glancing slices in small (<200 µm^2^) and large (>600 µm^2^) cells. The mean and standard deviation are shown. p > 0.05 by student’s t-test (n = 38 small and 70 large cells). Scale bar, 2 µm. (**E**) Maximum intensity projection of a confocal time series of a cell expressing ER marker Elo3-GFP (greyscale) and cytosol marker 3xmCherry (not shown) (DLY27372). Scale bar, 5 µm. (**F**) Quantification of cell volume (left), total ER content (middle), and ER density (right) for three representative cells. The top row is measured from the same cell shown in E. Red arrowheads indicate the times when mitosis took place. Grey boxes indicate the budded intervals.

**Supplemental Figure 2:**
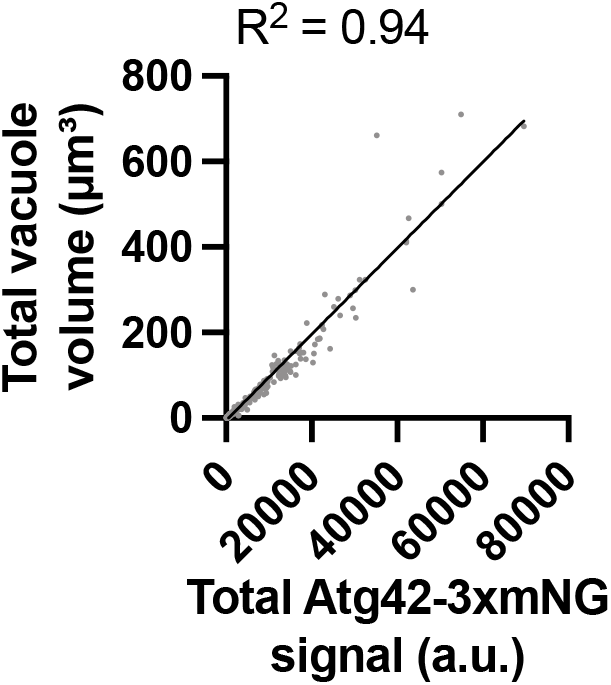
Total Atg42-3xmNG signal is correlated with vacuole volume. Plot of total Atg42-3xmNG signal and total vacuole volume per cell measured from cells expressing the vacuole lumen marker, Atg42-3xmNG, and cytosol marker, 3xmCherry (DLY25963). The best fit line, slope, and R^2^ values are shown (n = 375 cells).

**Supplemental Table 1.**

| Strain Name | Relevant Genotype | Source |
| --- | --- | --- |
| DLY24944 | *URA3:3xmCherry; SEC61-GFP:HYG<sup>R</sup>* | Wirshing et al., 2026 |
| DLY24947 | *URA3:3xmCherry; VPH1-GFP:HYG<sup>R</sup>* | Wirshing et al., 2026 |
| DLY25001 | *URA3:3xmCherry; PEX3-GFP:HYG<sup>R</sup>* | Wirshing et al., 2026 |
| DLY25312 | *URA3:3xmCherry; CIT1-GFP:NAT<sup>R</sup>* | Wirshing et al., 2026 |
| DLY25963 | *URA3:3xmCherry; ATG42-3xmNG:HYG<sup>R</sup>* | Wirshing et al., 2026 |
| DLY27372 | *URA3:3xmCherry; ELO3-GFP:NAT<sup>R</sup>* | Wirshing et al., 2026 |
| DLY27440 | *SEC61-3xmCherry:HYG<sup>R</sup>; Elo3-GFP:NAT<sup>R</sup>* | This study |

